# Anisotropy links cell shapes to a solid-to-fluid transition during convergent extension

**DOI:** 10.1101/781492

**Authors:** Xun Wang, Matthias Merkel, Leo B. Sutter, Gonca Erdemci-Tandogan, M. Lisa Manning, Karen E. Kasza

**Affiliations:** Department of Mechanical Engineering, Columbia University, New York, New York 10027, USA; Department of Physics, Syracuse University, Syracuse, New York, 13244, USA; Aix Marseille Univ, Université de Toulon, CNRS, CPT, Turing Center for Living Systems, Marseille, France

## Abstract

Within developing embryos, tissues flow and reorganize dramatically on timescales as short as minutes. This includes epithelial tissues, which often narrow and elongate in convergent extension movements due to anisotropies in external forces or in internal cell-generated forces. However, the mechanisms that allow or prevent tissue reorganization, especially in the presence of strongly anisotropic forces, remain unclear. We study this question in the converging and extending *Drosophila* germband epithelium, which displays planar polarized myosin II and experiences anisotropic forces from neighboring tissues, and we show that in contrast to isotropic tissues, cell shape alone is not sufficient to predict the onset of rapid cell rearrangement. From theoretical considerations and vertex model simulations, we predict that in anisotropic tissues two experimentally accessible metrics of cell patterns—the cell shape index and a cell alignment index—are required to determine whether an anisotropic tissue is in a solid-like or fluid-like state. We show that changes in cell shape and alignment over time in the *Drosophila* germband indicate a solid-to-fluid transition that corresponds to the onset of cell rearrangement and convergent extension in wild-type embryos and are also consistent with more solid-like behavior in *bnt* mutant embryos. Thus, the onset of cell rearrangement in the germband can be predicted by a combination of cell shape and alignment. These findings suggest that convergent extension is associated with a transition to more fluid-like tissue behavior, which may help accommodate tissue shape changes during rapid developmental events.

## Introduction

The ability of tissues to physically change shape and move is essential to fundamental morphogenetic processes that produce the diverse shapes and structures of tissues in multicellular organisms during development (1, 2). Developing tissues are composed of cells that can dynamically change their behavior and actively generate forces to influence tissue reorganization and movement (3–8). Remarkably, tissues dramatically deform and flow on timescales as short as minutes or as long as days (6). Recent studies highlight that tissue movements within developing embryos can be linked with the tissue fluidity (8–11). Fluid-like tissues accommodate tissue flow and remodeling, while solid-like tissues resist flow. Yet, the mechanisms underlying the mechanical behavior of developing tissues remain poorly understood, in part due to the challenges of sophisticated mechanical measurements inside embryos and the lack of unifying theoretical frameworks for the mechanics of multicellular tissues (6, 7, 12).

Epithelial tissue sheets play pivotal roles in physically shaping the embryos of many organisms (2), often through convergent extension movements that narrow and elongate tissues. Convergent-extension is highly conserved and used in elongating tissues, tubular organs, and overall body shapes (13). Convergent extension movements require anisotropies in either external forces that deform the tissue or asymmetries in cell behaviors that internally drive tissue shape change. Indeed, an essential feature of many epithelia *in vivo* is anisotropy in the plane of the tissue sheet, a property known as *planar polarity*, which is associated with the asymmetric localization of key molecules inside cells (14–17). For example, during *Drosophila* body axis elongation, the force-generating motor protein myosin II is specifically enriched at cell edges in the epithelial germband tissue that are oriented perpendicular to the head-to-tail body axis (18, 19) (Fig. 1*A*). Planar polarized myosin is required for cell rearrangements that converge and extend the tissue to rapidly elongate the body and is thought to produce anisotropic tensions in the tissue (19–25). In addition, the *Drosophila* germband experiences external forces from neighboring tissues, including the mesoderm and endoderm, which have been linked to cell shape changes in the germband during convergent extension (26–28) (Fig. 1*A*). Despite being fundamental to epithelial tissue behavior *in vivo*, it is unclear how such anisotropies arising from internal myosin planar polarity and external forces influence epithelial tissue mechanical behavior, particularly whether the tissue behaves more like a fluid or a solid.

**Figure 1.**
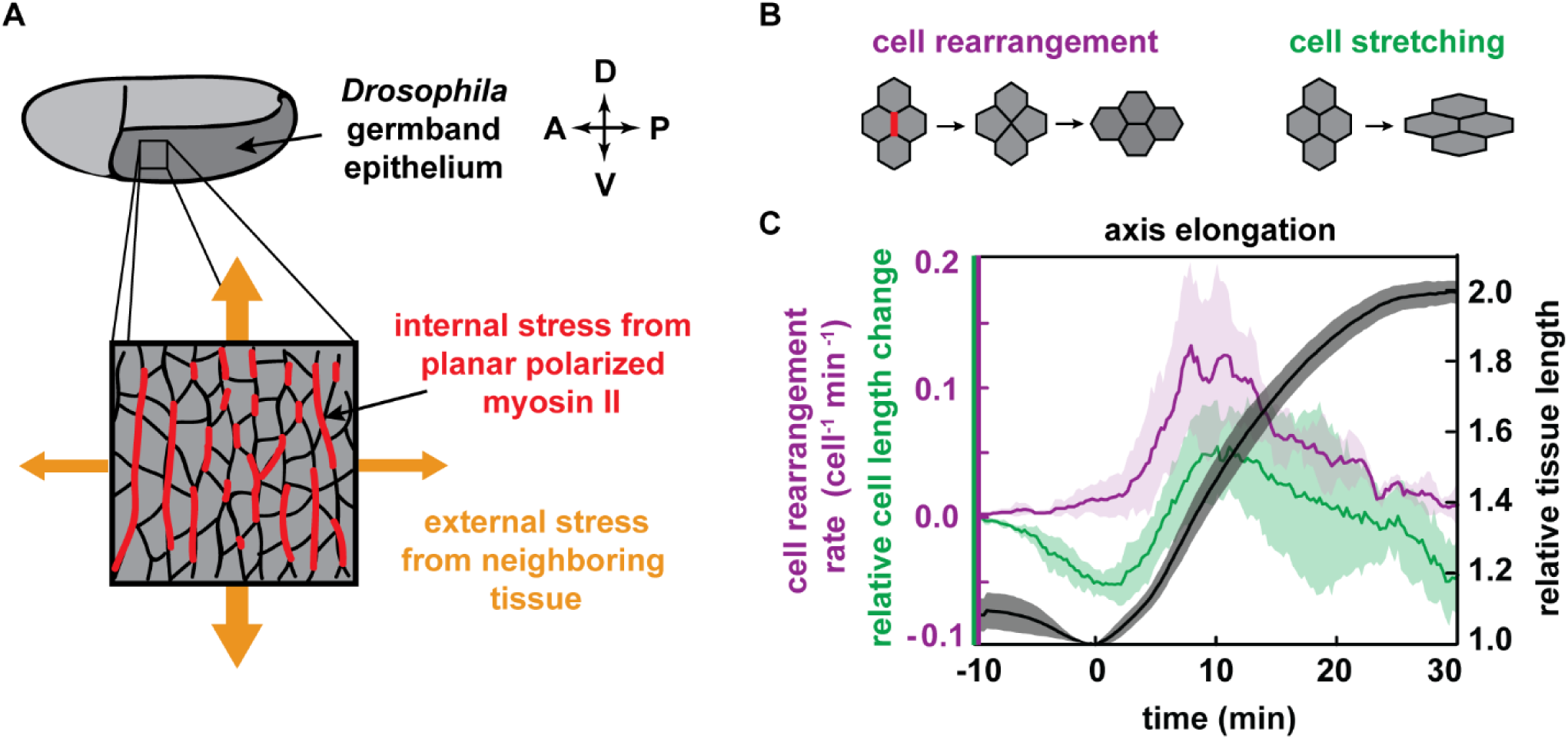
Cell shapes and cell rearrangements in the converging and extending *Drosophila* germband epithelium during axis elongation. (*A*) Schematic of *Drosophila* body axis elongation. The germband epithelium (dark gray) narrows and elongates along the head-to-tail body axis in a convergent extension movement. The tissue is anisotropic, experiencing internal stresses from planar polarized patterns of myosin II (red) within the tissue as well as external stresses (orange) due to the movements of neighboring tissue. (*B*) Schematic of oriented cell rearrangement and cell shape change. (*C*) The germband epithelium doubles in length along the head-to-tail axis in 30 min (black). Cell rearrangements are thought to drive tissue elongation (magenta), and cell shape changes also contribute (green). Tissue elongation begins at *t* = 0 . Relative cell length is normalized by the value at *t* = −10 min. Mean and standard deviation between embryos is plotted (*N*=5 embryos with an average of 278 cells per embryo per time point).

Vertex models have proven a useful framework for theoretically studying the mechanical behavior of confluent epithelial tissues (29, 30), including the packings of cells in tissues (31–33) and the dynamics of remodeling tissues (21, 31, 34, 35). Recent studies of the energy barriers to cell rearrangement in isotropic vertex models, which assume no anisotropy in either internal tensions at cell-cell contacts or in external forces, have revealed a transition from solid to fluid behavior, which depends on whether large or small contacts are favored between neighboring cells. The transition is indicated by a single parameter describing cell shape, 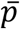, which is the average value in the tissue of cell perimeter divided by the square root of cell area (36–38). For small 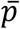, the tissue is solid-like, and above a critical value, 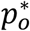, the tissue becomes fluid-like. The critical cell shape index 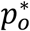 describes the shapes of cells at the solid-fluid transition. The precise value of 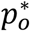 depends on the nature of cell packing in the tissue, and has a lower bound value of 3.81 (37, 39). The isotropic vertex model successfully predicts cell shapes at the transition from fluid-like to solid-like behavior in cultured primary bronchial epithelial tissues (39). Such a simple way to infer tissue behavior from static images is appealing, particularly for tissues that are inaccessible to mechanical measurements or live imaging. However, these previous vertex model studies did not account for effects of anisotropy, potentially limiting their use in the study of converging and extending tissues.

Here, we combine confocal imaging and quantitative image analysis with a vertex model of anisotropic tissues to study epithelial convergent extension during *Drosophila* body axis elongation. We show that cell shape alone is not sufficient to predict the onset of rapid cell rearrangement during convergent extension in the *Drosophila* germband, which exhibits anisotropies arising from internal forces from planar polarized myosin and external forces from neighboring tissue movements. Instead, we show that for anisotropic tissues, such as the *Drosophila* germband, anisotropy shifts the predicted transition from solid-like to fluid-like behavior and so must be taken into account, which can be achieved by considering both cell shape and cell alignment in the tissue. We find that the onset of cell rearrangement and tissue flow during convergent extension in wild-type and mutant *Drosophila* embryos is more accurately described by a combination of cell shape and alignment than by cell shape alone. These findings suggest that convergent extension is associated with a transition from solid-like to more fluid-like tissue behavior, which may help to accommodate dramatic epithelial tissue shape changes during rapid axis elongation.

## Results

### Cell shape alone is not sufficient to predict solid-like vs. fluid-like behavior in the *Drosophila* germband epithelium

To explore the mechanical behavior of a converging and extending epithelial tissue *in vivo*, we investigated the *Drosophila* germband, a well-studied tissue that has internal anisotropies arising from planar polarized myosin (18–23, 40) and also experiences external forces from neighboring developmental processes that stretch the tissue (26, 27). The germband rapidly extends along the anterior-posterior (AP) axis while narrowing along the dorsal-ventral (DV) axis (Fig. 1*A*), roughly doubling the length of the head-to-tail body axis in just 30 min (41) (Fig. 1*C*). Convergent extension in the *Drosophila* germband is driven by a combination of cell rearrangements and cell shape changes (Fig. 1*B,C*). The dominant contribution is from cell rearrangement (19, 20, 28, 41), which requires a planar polarized pattern of myosin localization across the tissue (18, 19) that is thought to be the driving force for rearrangement (19, 21, 22, 40). Cell stretching along the AP axis also contributes to tissue elongation and coincides with movements of neighboring tissues (26–28, 42, 43), indicating that external forces play an important role in tissue behavior. Despite significant study of this tissue, a comprehensive framework for understanding its mechanical behavior is lacking, in part because direct mechanical measurements inside the *Drosophila* embryo, and more generally for epithelial tissues *in vivo*, continue to be a challenge (44–46).

To gain insight into the origins of mechanical behavior in the *Drosophila* germband epithelium, we first tested the theoretical prediction of the vertex model that cell shapes can be linked to tissue mechanics. In the isotropic vertex model, tissue mechanical behavior is reflected in a single parameter, the average cell shape index 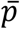 (36–39). To quantify cell shapes in the *Drosophila* germband, we used confocal time-lapse imaging of embryos with fluorescently-tagged cell membranes (47) and segmented the resulting time-lapse movies (28) (Fig. 2*A*). Prior to the onset of tissue elongation, individual cells take on roughly isotropic shapes and become more elongated over time (Fig. 2*A*,*B)*, consistent with previous observations (26–28, 48). Ten minutes prior to tissue elongation, the average cell shape index 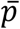 is just above 3.81. Eight minutes before the onset of tissue elongation 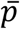 starts to increase before reaching a steady value of 3.98 about 20 min after the onset of tissue elongation (Fig. 2*B*). Remarkably, the average cell shape index in the tissue prior to tissue elongation, 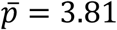, is close to the lower bound value predicted for the solid-fluid transition in the isotropic vertex model (dashed line in Fig. 2*B*) (36–38). This suggests that the tissue is solid-like prior to the onset of tissue elongation.

**Figure 2.**
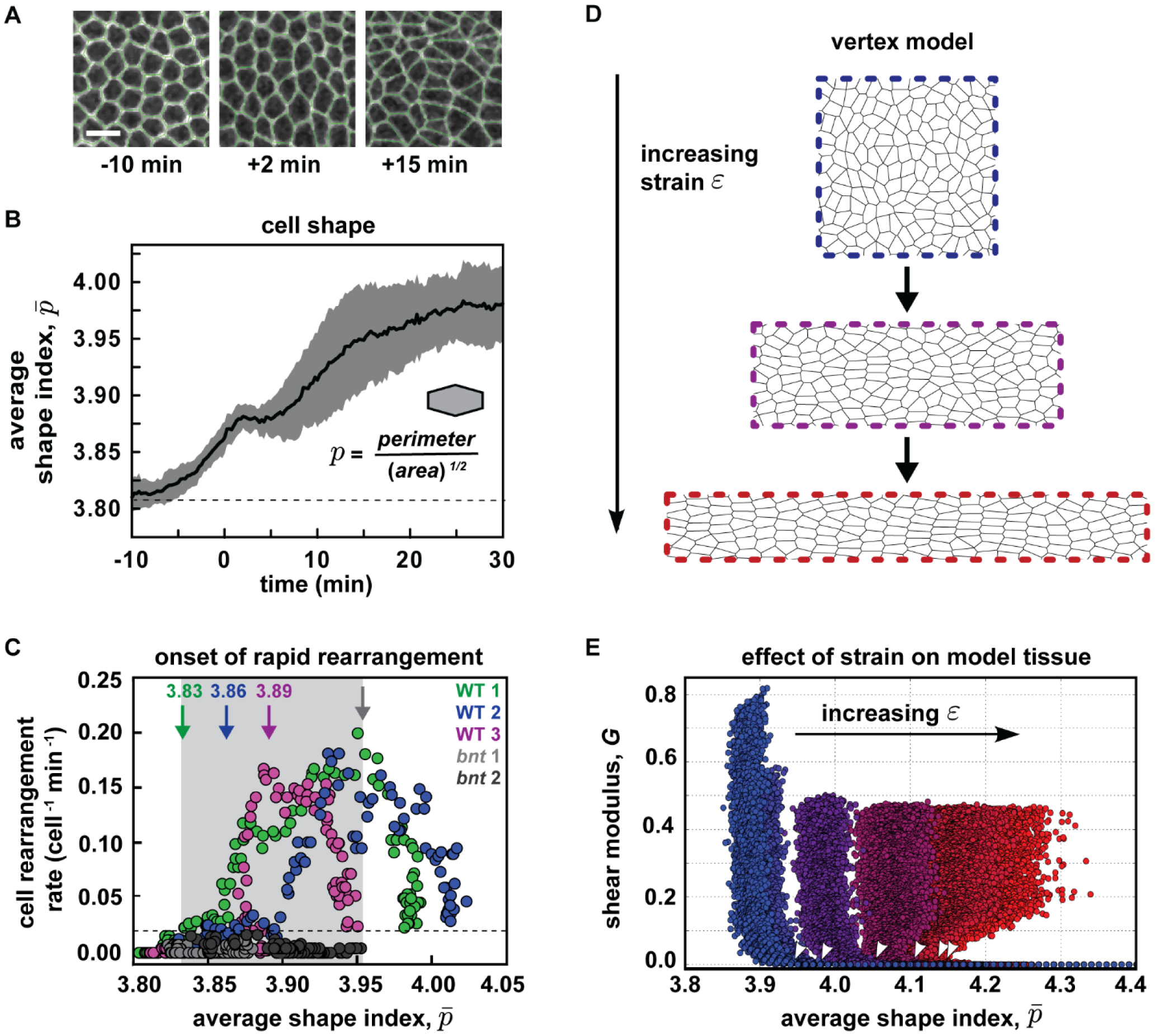
Cell shape alone is not sufficient to predict the mechanical behavior of anisotropic tissues such as the *Drosophila* germband. (*A*) Confocal images from time lapse movies of epithelial cell patterns in the ventrolateral region of the germband tissue during *Drosophila* body axis elongation. Cell outlines were visualized using the fluorescently-tagged cell membrane marker, gap43:mCherry (47). Anterior left, ventral down. Images of cells with overlaid polygon representations used to quantify cell shapes. (*B*) The average cell shape index 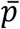 in the germband tissue before and during convergent extension. The cell shape index, *p*, is calculated for each cell from the ratio of cell perimeter to the square root of cell area, and the average value for cells in the tissue, 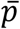, is calculated at each time point. The mean and standard deviation between embryos is plotted. Dashed line denotes the lower bound value for the solid-to-fluid transition in the isotropic vertex model, 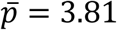. (*C*) The instantaneous rate of cell rearrangements per cell versus the average cell shape index 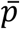 from movies of individual *Drosophila* embryos at time points before and during convergent extension (3 wild-type embryos and 2 mutant *bnt* embryos). Green, blue, and magenta arrows indicate the values of 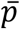 at the onset of rapid cell rearrangement (>0.02 per cell per min, dashed line) in three different wild-type embryos. Gray arrow indicates the highest measured value of 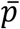 in a mutant *bnt* embryo, and for which rapid cell rearrangement in the tissue had still not occurred. Shaded gray region denotes values of 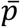 for which different embryos display distinct mechanical behaviors, either showing rapid cell rearrangement or remaining largely static. Thus, 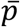 alone is not sufficient to determine the onset of rapid cell rearrangement in the tissue. (*D*) We study the effect of anisotropies on the solid-fluid transition in the vertex model by externally applying an anisotropic strain *ε*. An initially quadratic periodic box with dimensions *L*_0_ × *L*_0_ is deformed into a box with dimensions *e*^*ε*^*L*_0_ × *e*^−*ε*^*L*_0_. (*E*) Vertex model tissue rigidity as a function of the average cell shape index with different levels of externally applied strain *ε* (values for *ε*, increasing from blue to red: 0, 0.2, 0.4, 0.8, 1). For every force-balanced configuration, the shear modulus was analytically computed as described in the Supplementary Materials and Methods. For zero strain, we find a transition at an average cell shape index of 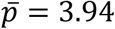 from rigid, solid-like behavior to floppy, fluid-like behavior. See also Fig. S1 for effects of cell packing disorder on the transition. For increasing strain, the transition from solid-like to fluid-like behavior (i.e. the shear modulus becomes zero for a given strain) occurs at higher 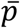 (approximate positions marked by white arrows). Thus, a single critical cell shape index is not sufficient to determine the transition from rigid, solid-like to floppy, fluid-like behavior in an anisotropic tissue.

We next asked how these cell shapes correlate with tissue mechanical behavior. As an experimentally accessible read-out of tissue fluidity, we used the instantaneous rate of cell rearrangements occurring within the germband tissue (Fig. 1*C*, magenta), with higher rearrangement rates associated with more fluid-like behavior and/or larger active driving forces. Plotting instantaneous cell rearrangement rate versus 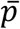 at each time point from movies of individual wild-type *Drosophila* embryos, we find that the onset of rapid cell rearrangement occurs at different values of 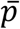 for each embryo, ranging from 3.83 to 3.89 for a threshold rearrangement rate per cell of 0.02 min^-1^ (Fig. 2*C*). Moreover, in *bcd nos tsl* (*bnt*) mutant embryos, which lack anterior-posterior patterning genes required for axis elongation, 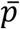 reaches a value of 3.95 without the tissue undergoing significant cell rearrangement (Fig. 2*C*, gray symbols). By comparison, cells in wild-type embryos undergo rapid cell rearrangement for this value of 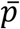 Thus, in the germband epithelium, the cell shape index 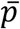 alone is not sufficient to predict tissue behavior.

### Cell shape alone is not sufficient to predict solid vs. fluid behavior in a vertex model of anisotropic tissues

The result that cell shape alone is not sufficient to predict tissue behavior seems to contradict the previous predictions in the isotropic vertex model. To study whether anisotropies in the germband could also play a role, we used vertex model simulations to test how tissue anisotropy, introduced into the model in two different ways, affects the predicted behavior. First, we introduced anisotropy by applying an external deformation, mimicking the effects of forces exerted by neighboring morphogenetic processes, and then studying force-balanced states of the model tissue (Fig. 2*D*). As a metric for tissue solidity (as opposed to fluidity), we measured the shear modulus of the model tissue, which reflects tissue rigidity or equivalently how much the tissue resists changes in shape. We then analyzed how the shear modulus correlates with the average cell shape index 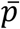 for different values of the deforming strain *ε* (Fig. 2*E*).

For small deformations (small strain *ε*), we recover the behavior of the isotropic vertex model. The shear modulus is finite (solid-like behavior) when 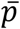 is small and vanishes (fluid-like behavior) above a critical cell shape index, which is 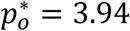 for our simulations (Fig. 2*E*, blue symbols). This is above the previously reported 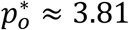 (36, 37), but it has previously been noted that 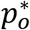 should depend on exactly how cells are packed in the tissue (49–51), a property referred to as the packing disorder. The packing disorder is non-existent for a cellular packing consisting only of hexagons. In contrast, the existence of cells with a neighbor number different from six at random places will increase the packing disorder. We performed a large number of vertex model simulations to confirm that the critical 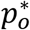 depends on the packing disorder by systematically varying the *in silico* preparation protocol (Fig. S1*A,B*), which suggests that in experiments 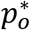 can be treated as a fitting parameter that roughly characterizes the packing disorder in a tissue.

For larger deformations, we study how the critical value of the shape index at the transition between solid-like and fluid-like behavior changes with the amount of strain. We find that it generally increases with strain (Fig. 2*E*). Indeed, 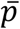 for cells in a deformed, solid-like tissue (with positive shear modulus) can be higher than for cells in an undeformed, fluid-like tissue (with vanishing shear modulus). Thus, consistent with our observations of the *Drosophila* germband, the cell shape index alone is not sufficient to predict mechanical behavior in an anisotropic tissue.

### Theoretical considerations and vertex model simulations predict a shift of the solid-fluid transition in anisotropic tissues

Some of us recently developed a theoretical understanding for a shift in the critical shape index when deforming a vertex model tissue (51). In the limit of small deformations by some strain *ε* and without cell rearrangements, the critical value of 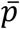 increases from 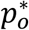 to 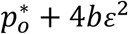, where *b* is a constant prefactor. To compare this formula to the vertex model simulations (Fig. 2*D,E*), we need to take into account that cell rearrangements occur in our simulations. Removing their contribution from the overall tissue strain *ε* leaves us with a parameter *Q* (Fig. 3*A* and Supplementary Materials and Methods), which can be quantified using a triangulation of the tissue created from the positions of cell centers (52, 53). We term *Q* a “shape alignment index”, as *Q* is non-zero only when the long axes of cells are aligned. We emphasize that, unlike the nematic order parameter for liquid crystals, the cell alignment parameter *Q* is additionally modulated by the degree of cell shape anisotropy; tissues with the same degree of cell alignment but more elongated cells have a higher *Q* (Fig. 3*A*). In other words, *Q* can be regarded as a measure for tissue anisotropy. After accounting for cell rearrangements, we expect the transition point in anisotropic tissues to shift from the isotropic value 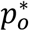 to (Supplementary Materials and Methods):

**Figure 3.**
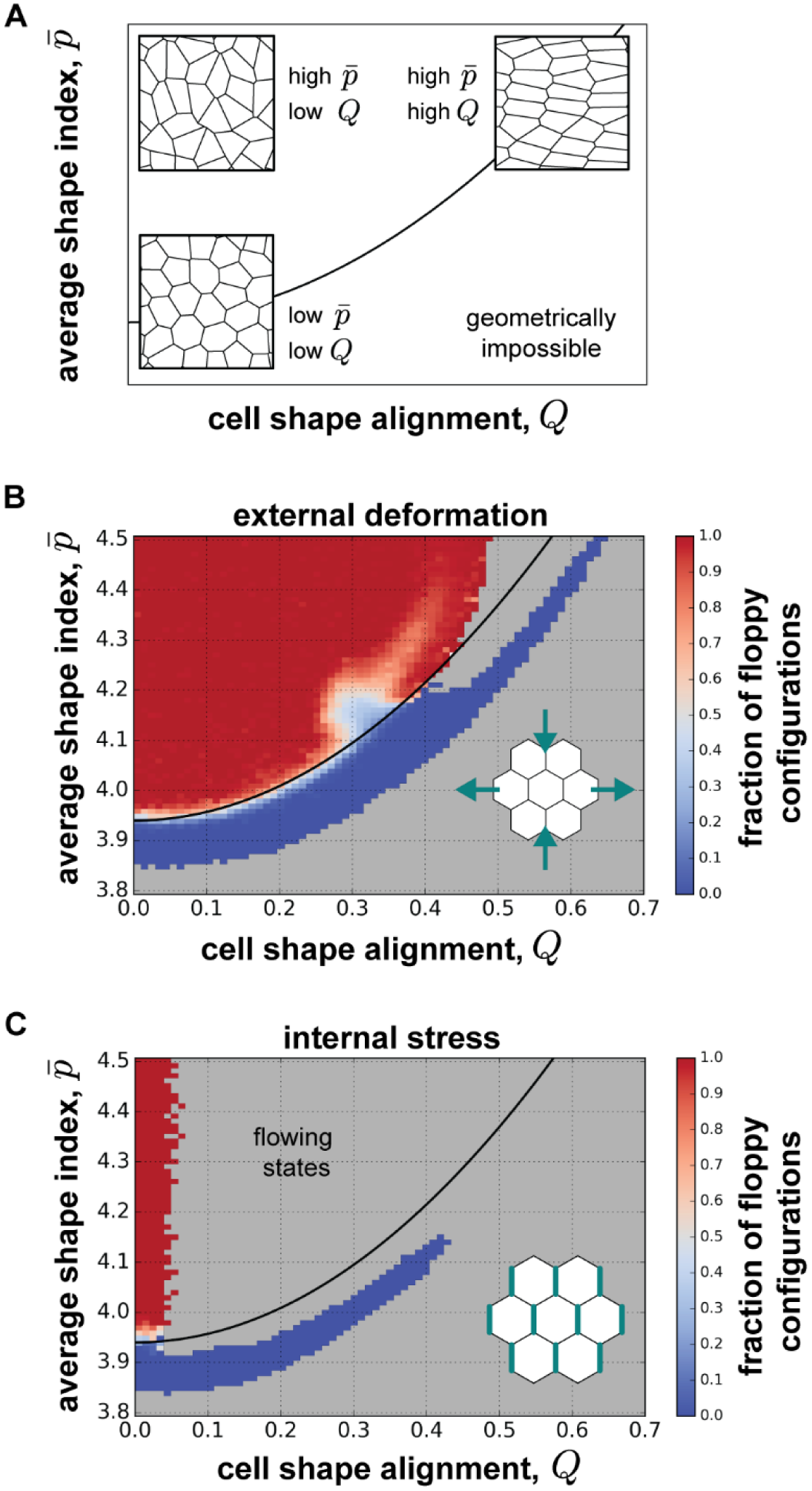
The solid-to-fluid transition in a vertex model of anisotropic tissues. (*A*) Cell shape and cell shape alignment can be used to characterize cell patterns in anisotropic tissues. Cell shape alignment *Q* characterizes both cell shape anisotropy and cell shape alignment across the tissue. While a high cell shape index 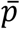 correlates with anisotropic cell shapes, the cell shape alignment *Q* is only high if these cells are also aligned. Conversely, low 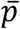 implies low cell shape anisotropy and thus low *Q*. (*B-C*) Vertex model simulations for the case of an anisotropic tissue arising (*B*) due to externally induced deformation (cf. Fig. 2*D,E*) or (*C*) due to internal active stresses generated by cells. The fraction of fluid-like, floppy configurations is plotted as a function of the average cell shape index 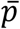 and cell shape alignment *Q*. For both internal and external sources of anisotropy, the critical shape index 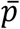 marking the transition between solid-like, rigid states (blue) and fluid-like, floppy states (red) is predicted to depend quadratically on *Q*. Gray regions denote combinations of 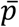 and *Q* for which we did not find force-balanced states. In panel *B*, the black solid line shows a fit of the transition to Eq. (1) with 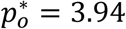 and *b* = 0.43, and panel *C* shows this same line. In panel *B*, a deviation from Eq. (1) is only seen around 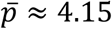 and *Q* ≈ 0.3, which is likely due to effects of packing disorder.

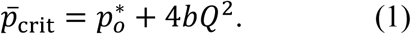

Indeed, comparing this equation to vertex model simulations yields a good fit with the simulation results (Fig. 3*B*, black solid line), with fit parameters 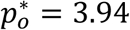 and *b* = 0.43. While we would generally expect the precise value of *b* to also depend on the packing disorder, our best-fit value is within the previously determined range (51). Hence, for external deformation, the solid-fluid transition point in the vertex model increases quadratically with tissue anisotropy *Q*.

We also tested how the model predictions change when we instead introduce anisotropy generated by internal forces into the vertex model. We modeled myosin planar polarity as increased tensions on “vertical” cell-cell contacts (21), and focus again on stationary, force-balanced states (Fig. 3*C*). Similar to the case of external deformation, we find that solid states exist for larger cell shape indices than the isotropic 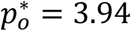, and our results are again consistent with the fit from Fig. 3*B* (black solid line). With finite anisotropic internal tensions we obtain permanently flowing states in the fluid-like (floppy) regime that do not reach a force-balanced state, and this explains the gray region devoid of stable states in the upper middle region of Fig. 3*C*. Taken together, these findings demonstrate that a combination of cell shape and cell shape alignment in the vertex model indicates whether an anisotropic tissue is in a solid-like or fluid-like state, regardless of the underlying origin of anisotropy.

### Cell shape and cell shape alignment together indicate a solid-to-fluid transition during *Drosophila* axis elongation that corresponds to the onset of cell rearrangement

We returned to our experiments to test whether a combination of 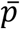 and *Q* would be a better predictor for the behavior of the *Drosophila* germband during convergent extension. We quantified alignment *Q* using the triangle method (Fig. 4*A*) and found that prior to the onset of tissue elongation, which begins at *t* = 0 min, alignment is not very high (Fig. 4*B*). *Q* begins to increase just prior to elongation, peaking at *t* = 1 min (Fig. 4*B*), which is consistent with observations using other cell pattern metrics (21, 26, 28). This peak in *Q* corresponds to stretching of the cells along the dorsal-ventral axis, perpendicular to the axis of germband extension, and coincides with the time period during which the presumptive mesoderm tissue is invaginating (47). We observe that *Q* relaxes back to low levels during axis elongation (Fig. 4*B*). Thus, cell shapes in the germband tissue are transiently aligned around the onset of convergent extension.

**Figure 4.**
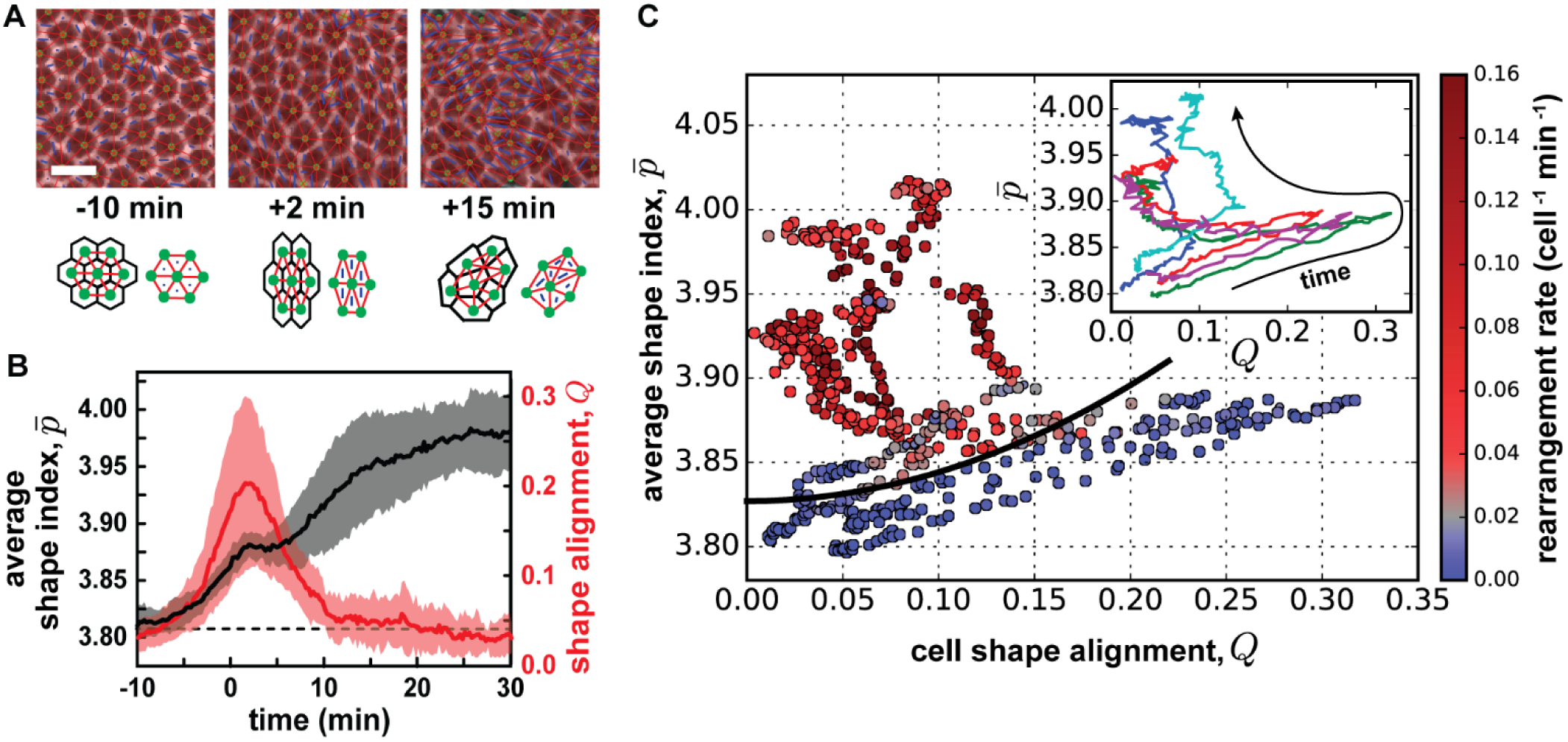
Cell shape and cell shape alignment together indicate a solid-to-fluid transition during *Drosophila* convergent extension that corresponds to the onset of cell rearrangement. (*A*) Confocal images from time lapse movies of epithelial cell patterns in the ventrolateral region of the germband tissue during *Drosophila* body axis elongation. Cell outlines were visualized using the fluorescently-tagged cell membrane marker, gap43:mCherry (47). Anterior left, ventral down. Scale bar, 10 μm. Images of cells with overlaid triangles that were used to quantify cell shape anisotropy. Cell centers (green dots) are connected with each other by a triangular network (red bonds). Cell shape stretches are represented by triangle stretches (blue bars), and the average cell elongation, *Q*, is measured (53). (*B*) The cell shape alignment index *Q* (red) and average cell shape index 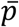 (black, same as Fig. 2*B*) for the germband tissue before and during axis elongation. The cell shape alignment index *Q* was calculated for each time point, and the mean and standard deviation between embryos is plotted (*N*=5 embryos with an average of 278 cells per embryo per time point). The onset of tissue elongation occurs at *t* = 0. Dashed line denotes the lower bound value for the solid-to-fluid transition in the isotropic vertex model, 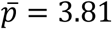. (*C*) The relationship between the average cell shape index 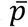 and cell shape alignment *Q* for five individual wild-type embryos, with each point representing 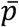 and *Q* for a single time point in a single embryo. Instantaneous cell rearrangement rate per cell in the tissue is represented by the color of each point, with blue indicating low rearrangement rate and red indicating high rearrangement rate. The black solid line indicates a fit to Eq. (1), from which we extract 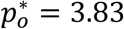, where *b* was fixed to the value obtained in vertex model simulations (cf. Fig. 3*B*). *Inset:* Each color represents an individual embryo. 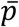 increases over time. *Q* increases nearly linearly with 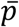, when 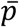 is less than 3.87, which corresponds to the time period prior to the onset of cell rearrangement. At the onset of cell rearrangement and tissue remodeling, *Q* drastically drops as 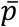 continues to increase.

We next asked whether this temporary increase in alignment could help resolve the seeming contradiction between the measured cell shapes and cell rearrangement rates. We compared our measurements of 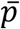 and *Q* before and during axis elongation. Plotting 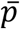 vs *Q* at each time point in movies of individual wild-type embryos reveals common features, despite embryo-to-embryo variability (Fig. 4*C*, inset). Initially, we see a concomitant increase of 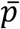 and *Q* prior to the onset of convergent extension. Above 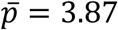, *Q* decreases drastically as 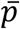 continues to increase, indicating that further increases in 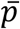 are associated with randomly oriented cell shapes (cf. Fig. 3*A*).

Next, we investigated how 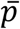 and *Q* correlate with the instantaneous rate of cell rearrangements occurring within the germband, with higher rearrangement rates associated with more fluid-like behavior and/or larger active driving forces (Fig. 4*C*). To isolate the changes in mechanical properties of the germband during convergent extension from later developmental events, we focus on times *t* ≤ 20 min after the onset of tissue elongation, well before cell divisions begin in the germband. The anisotropic vertex model predicts that the solid-like or fluid-like behavior of the tissue should depend on both 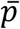 and *Q* according to Eq. (1), with only two adjustable parameters, 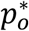 and *b*. We fit Eq. (1) to our experimental data by minimizing the number of experimental data points on the wrong side of the theoretical transition line, and for simplicity vary only 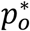 while keeping the theoretically determined value for *b*. To differentiate between solid-like and fluid-like tissue behavior in the experimental data, we choose a cutoff on the cell rearrangement rate of 0.02 min^-1^, and we find 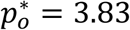 (Fig. 4*C*). While different cutoff values lead to differences in 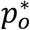, our general conclusions remain unchanged (Fig. S2*B*). During early times, when 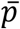 and *Q* are both increasing, the tissue stays within the predicted solid-like regime (Fig. 4*C*). The subsequent rapid decrease in *Q* brings embryos closer to the transition line. As cell shape indices further increase, individual embryos cross this transition line at different points, coinciding with increased rates of cell rearrangement. Thus, compared to the isotropic model, the anisotropic vertex model improves the prediction of the onset of rapid cell rearrangement and tissue flow during convergent extension with two metrics of cell patterns, 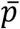 and *Q*, that are both easy to access experimentally.

### Cell shape and alignment help to explain tissue behavior in *bcd nos tsl* mutant embryos

Since the *Drosophila* germband experiences both internal forces due to myosin planar polarity and external forces from neighboring tissues, we wondered whether our theoretical predictions still hold when altering the nature of the forces in the germband. To dissect the effects of internal and external sources of tissue anisotropy, we studied cell patterns in *bcd nos tsl* (*bnt*) mutant embryos, which lack anterior-posterior patterning genes required for axis elongation. These mutant embryos do not display myosin planar polarity (Fig. 5*A*) and show severe defects in tissue elongation (Fig. 5*C*), cell rearrangement (Fig. 5*D*), and endoderm invagination, but still undergo mesoderm invagination (18, 20, 25, 26, 28, 41).

**Figure 5.**
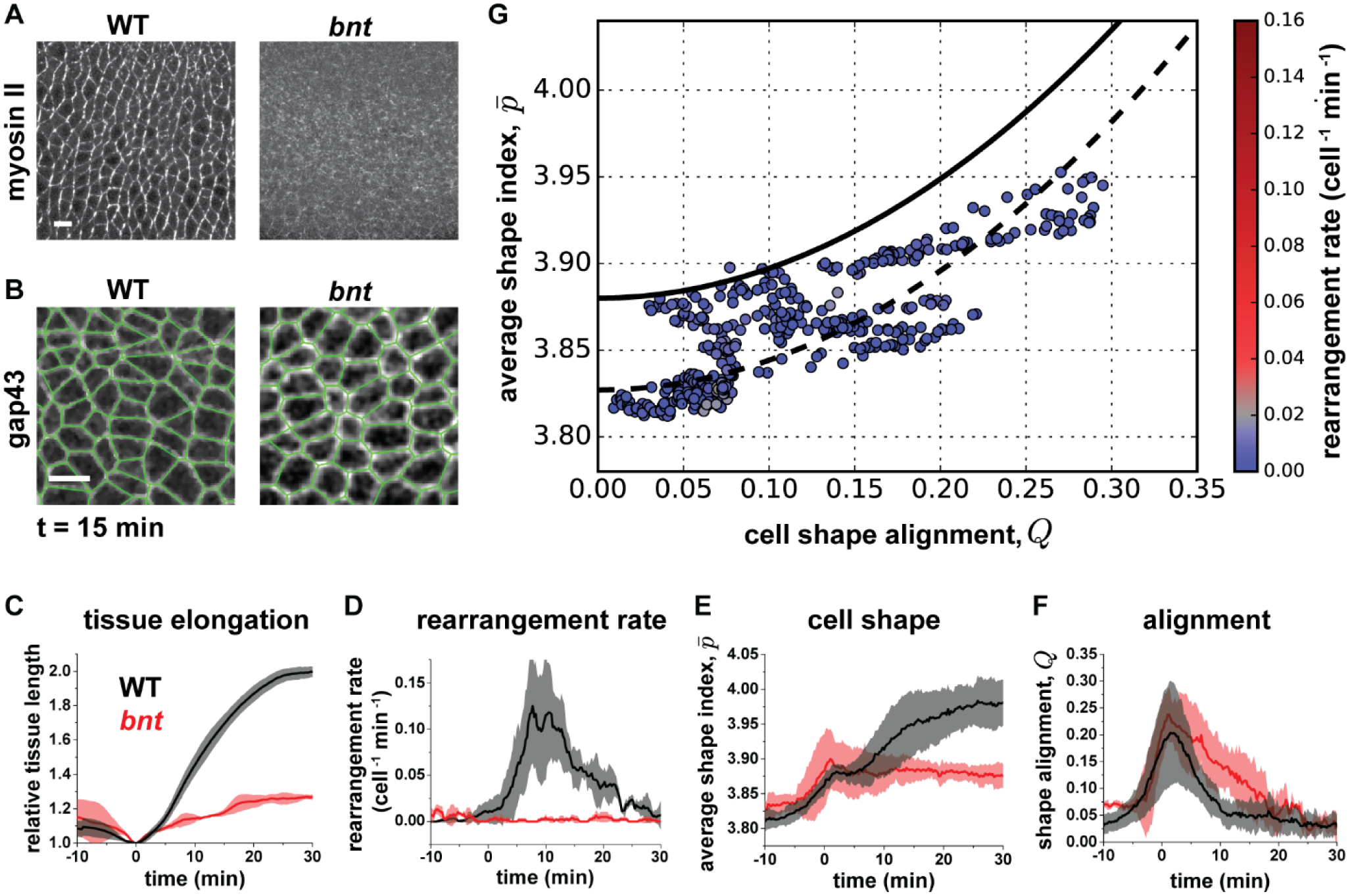
Cell shape and cell shape alignment help to explain the solid-like behavior of the germband in *bnt* mutant embryos. (*A*) *bcd nos tsl* (*bnt*) mutant embryos, which lack anterior-posterior patterning genes required for axis elongation, show severely disrupted myosin planar polarity compared to wild-type embryos. (*B*) Confocal images from time lapse movies of epithelial cell patterns at *t* = +15 min with outlines visualized using the fluorescently-tagged cell membrane marker, gap43:mCherry (47). Polygon representations of cell shapes used to quantify cell patterns are overlaid (green). Scale bars, 10 μm. (*C*) Tissue elongation is severely reduced in *bnt* mutant embryos compared to wild-type controls. (*D*) Cell rearrangement rate per cell in the tissue is severely reduced in *bnt* mutant embryos. (*E*) The average cell shape index 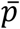 in the tissue shows similar behavior to wild-type controls for *t* < 5 min, but unlike in wild-type embryos, 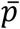 does not show further increases with time for *t* > 5 min. (*F*) The cell alignment index *Q* shows similar behavior to wild-type control embryos for *t* < 5 min, but *Q* relaxes more slowly to low levels in *bnt* compared to wild-type control embryos. (*C-F*) The mean and standard deviation between embryos is plotted (*N* = 3 *bnt* embryos with an average of 190 cells per embryo per time point). (*G*) The relationship between 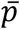 and *Q* for three individual *bnt* mutant embryos, with each point representing 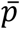 and *Q* for a single time point in a single embryo. Instantaneous cell rearrangement rate in the tissue is represented by the color of each point, with blue indicating low rearrangement rate and red indicating high rearrangement rate. We find a lower bound for 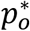 in Eq. (1) of 3.88 (black solid line), which leaves only two data points in the fluid regime that are likely outliers. This value is above the wild-type value (dashed line, cf. Fig. 4*C*) pointing to potentially different packing disorder in wild-type and *bnt* germband tissue.

In *bnt* embryos, the average cell shape index 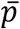 displays an initial increase until *t* = 1 min (Fig. 5*E*), concomitant with an increase in cell shape alignment *Q* (Fig. 5*F*). These observations are consistent with the idea that external forces from the mesoderm produce this initial increase in cell elongation and alignment, and are similar to our observations in control embryos, although with a somewhat larger cell shape index 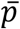 for the *bnt* case. However after *t* = 1 min in *bnt* embryos, 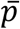 does not increase further and instead takes on a steady value of 3.87 during the rest of the process (Fig. 5*E*). This supports the idea that the further increase in 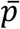 observed in wild-type embryos is due to internal anisotropies associated with myosin planar polarity or external forces associated with endoderm invagination. Interestingly, *Q* returns more slowly to low levels in *bnt* compared to control embryos (Fig. 5*F*), suggesting a potential role for myosin planar polarity, cell rearrangement, or endoderm invagination in relaxing cell shape alignment.

The increased cell shape index in *bnt* embryos around *t* = 1 min would have erroneously indicated more fluid-like behavior in a purely isotropic vertex model. Instead, plotting 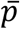 vs *Q* for each time point in individual *bnt* embryos reveals that the time points for which the tissue exhibits high 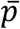 also exhibit high alignment *Q* (Fig. 5*G*). Since the *bnt* tissues do not transition to a state of rapid cell rearrangement and remain below our cutoff on rearrangement rate of 0.02 min^-1^, we cannot directly fit Eq. (1) to this data. However, we identified a lower bound of 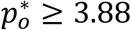 (Fig. 5*G*, black solid line). This suggests that 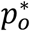 for the germband in *bnt* mutant embryos is likely higher than in wild-type embryos (Fig. 5*G*, black solid and dashed lines), raising the possibility that the precise value of 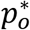 is genotype-specific due to differences in cell packing disorder. Thus, taking cell shape alignment *Q* into account can help to reconcile the solid-like tissue behavior in mutant *bnt* embryos with the increased shape index 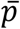, even in the absence of internal anisotropic tensions induced by myosin polarity.

## Discussion

In this work, we show that cell shape and alignment can be used to understand and predict whether an anisotropic tissue behaves more like a fluid that accommodates flow and remodeling or instead behaves like a solid that tends to maintain its shape. Importantly, we find that, in contrast to isotropic tissues, the mechanical behavior of the converging and extending *Drosophila* germband cannot be predicted by cell shape alone. Instead, we show via theoretical analysis and simulation that in anisotropic tissues two experimentally accessible metrics—the cell shape index complemented by a cell alignment index—are required to determine whether an anisotropic tissue is in a solid-like or fluid-like state, regardless of the underlying origin of anisotropy. We demonstrate that the onset of rapid cell rearrangement in wild-type *Drosophila* embryos is indeed more accurately described by a combination of these two cell pattern metrics than by cell shape alone. In addition, we find that a combination of cell shape and alignment help to explain the more solid-like behavior in the germband of *bnt* mutant embryos. These findings suggest that convergent extension of the *Drosophila* germband can be viewed as a transition from solid-like to more fluid-like mechanical behavior to help accommodate dramatic tissue shape changes and are consistent with the idea that the properties of developing tissues can be tuned to become more fluid-like during rapid morphogenetic events.

We present a theoretical framework for predicting the mechanical behavior of anisotropic epithelial tissues with two metrics describing cell patterns, and we demonstrate that these metrics predict the onset of rapid cell rearrangement in the *Drosophila* germband. There are only two adjustable parameters in the model, 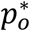 and *b*, which capture the cell shape index at the solid-fluid transition in an isotropic tissue and the strength of tissue anisotropy effects, respectively. Effects of cell alignment characterized by *b* are required to understand the mechanical behavior of the wild-type germband. Meanwhile, effects of packing disorder can generally affect the precise value of both 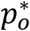 and *b*. Such effects may explain a general shift of the isotropic transition point 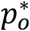 between wild type and *bnt* embryos. Future theoretical studies will uncover how 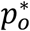 and *b* depend on cell packing disorder. Remarkably, the vertex model predictions of tissue behavior are independent of the underlying origin of anisotropy, and therefore can be used to predict mechanical properties of tissues from cell shape patterns, even when external stresses and internal active stresses cannot be directly measured. Importantly, the average cell shape index 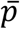 and cell shape alignment index *Q* are easy to access experimentally from snapshots of cell packings in tissues, even in systems where time-lapse live imaging of cell rearrangement and tissue flow is not possible. Thus, this approach may prove useful for studying complex tissue behaviors in a broad range of morphogenetic processes occurring in developing embryos *in vivo* or organoid systems *in vitro*.

Our result of a higher isotropic transition point 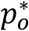 in mutant *bnt* embryos than in wild-type embryos suggests that due to differences in cellular packing disorder, the barriers to cell rearrangement and tissue remodeling are higher in mutant *bnt* embryos. The *Drosophila* germband epithelium displays internal anisotropies from myosin planar polarity and also experiences external forces from neighboring tissues, including the invaginating mesoderm and endoderm, whereas *bnt* embryos experience altered forces associated with reduced myosin planar polarity and defects in endoderm invagination. While this alone might be sufficient to explain axis elongation defects observed in *bnt* mutant embryos, higher barriers to cell rearrangement would contribute to the elongation defects. Moving forward, it will be interesting to explore experimentally how packing affects barriers to cell rearrangement in different tissues, how the nature of fluctuations driven by actomyosin contractility, cell tractions, or cell proliferation during tissue formation contribute to these barriers, and how these factors impact tissue mechanics and remodeling during development and in disease states. How internal and external forces together influence myosin localization and dynamics, cell rearrangement, cell shape change, and tissue flow remain open questions, and the approaches we develop here will be useful for interrogating these questions.

Prior to the onset of cell rearrangement and tissue flow, cells in the *Drosophila* germband take on shapes near the lower bound value for the solid-fluid transition predicted by the isotropic vertex model. This suggests that the initially solid-like tissue may be poised near this transition to allow for rapid cell rearrangement and tissue flow with small changes in cell properties or under small applied forces, while still maintaining tissue integrity. This is possibly consistent with recent ferrofluid droplet and magnetic bead microrheology measurements of tissues in the cellularizing *Drosophila* embryo prior to axis elongation, which indicated that tissue behavior is predominantly elastic (solid-like) over timescales less than several minutes (45, 46). While both studies compared the long-time tissue behavior to fluid-like viscous models, their observations might also be consistent with a weak yield-stress solid, which is solid-like for small stresses but deforms plastically and flows for larger stresses. This interpretation would be supported by the near absence of cell rearrangements during this developmental stage.

During axis elongation in *Drosophila*, we find that cell shape and cell alignment in the germband change in a manner predicted to modulate the solid-like or fluid-like behavior of the tissue. However, we note that the tissue does not deviate far from the predicted solid-fluid transition line. In addition, cell rearrangements are not randomly oriented. The large majority of rearrangements are oriented along the head-to-tail body axis (19, 20, 40, 41, 54), and the time period of rapid cell rearrangement (Fig. 1*C*) coincides with the period of planar polarized myosin (23, 25, 40). Taken together, these findings suggest that over the 20 min timescale of rapid axis elongation, the *Drosophila* germband tissue may still behave as a weak yield-stress solid, in which driving forces from planar polarized myosin are required to overcome energy barriers to drive cell rearrangements and tissue flow over these short timescales.

After 20 minutes of axis elongation, cell rearrangement rates decrease back to low levels (Fig. 1*C*). This slowing of rearrangement during axis elongation is not well understood. It may be related to changes in myosin organization and dynamics in germband cells preparing for cell divisions that begin around *t* = 30 min (55). Alternately, this may be related to changes in cellular adhesive interactions over the course of embryonic development (56, 57). Interestingly, the anisotropic vertex model presented here would predict fluid-like behavior for the tissue during this period of slowing cell rearrangement, given the observed high values of cell shape index 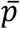 and low values of cell alignment index *Q* (Fig. 4*B,C*). One possible explanation for the discrepancy is that driving forces are no longer sufficient to produce rapid cell rearrangement. Consistent with this possibility, myosin planar polarity reaches a maximum 5-10 min after the onset of axis elongation and then decreases during the rest of the process (23, 28, 40). Alternatively, the description of cell rearrangements in the vertex model may be too simple for epithelial tissues with more mature cell-cell junctions, where additional barriers to cell rearrangements may occur. Future studies will be needed to explore these possibilities and incorporate biologically sophisticated pictures of cell rearrangement into models.

Interestingly, a fluid-to-solid jamming transition has recently been reported in mesodermal tissues during zebrafish body axis elongation (8). In contrast to the zebrafish mesoderm in which the transition to more solid-like behavior is associated with an increase in cellular volume fraction (proportion of the tissue occupied by cells), the *Drosophila* germband epithelium comprises tightly packed cells and its mechanical behavior changes in the absence of any change in cell volume fraction. Future studies will be needed to explore how the contractile and adhesive properties of epithelial cells might be regulated during development to tune the solid-like or fluid-like mechanical behaviors of the tissues in which they reside.

## Methods

Embryos were generated at 23°C and analyzed at room temperature. Cell outlines were visualized with gap43:mCherry (47) or Resille:GFP cell membrane markers. Embryos were imaged on a Zeiss LSM880 laser scanning confocal microscope. Time-lapse movies were analyzed with SEGGA software in MATLAB (28) for quantifying cell shapes and cell rearrangement rates, PIVlab version 1.41 in MATLAB (58) for quantifying tissue elongation, and custom MATLAB code for quantifying cell alignment using the triangle method (52, 53, 59). The vertex model describes an epithelial tissue as a planar tiling of *N* cellular polygons, where the degrees of freedom are the vertex positions (32). Forces in the model are defined such that cell perimeters and areas act as effective springs with a preferred perimeter *p*_0_ and a preferred area of one, which is implemented via an effective energy functional (51). Unless otherwise noted, error bars are the standard deviation. Details can be found in the Supplementary Materials and Methods.

## Acknowledgements

The authors thank Erik Boyle for assistance with data processing; Dene Farrell and Jennifer Zallen for the use of SEGGA, a segmentation and quantitative image analysis toolset (28); Adam Martin for the sqh-gap43:mCherry fly stock (47); and the Bloomington Drosophila Stock Center (BDSC) for fly stocks. This work was supported by the National Science Foundation CMMI 1751841 to K.E.K., DMR-1352184 and POLS-1607416 to M.L.M, and DMR-1460784 (REU) to L.B.S. M.L.M., M.M., and G.E.T. acknowledge support from Simons Grant No. 446222 and 454947, and NIH R01GM117598. K.E.K. holds a BWF Career Award at the Scientific Interface, Clare Boothe Luce Professorship, and Packard Fellowship.

