## Supplementary Information for "Anisotropy links cell shapes to a solid-to-fluid transition during convergent extension"

### Supplementary Materials and Methods

**Fly Stocks and Genetics.** Embryos were generated at 23°C and analyzed at room temperature. Wild-type control embryos were *yw* with one maternal copy of a sqh-gap43:mCherry transgene to label cell membranes (1). The *bcd nos tsl* (*bnt*) maternal mutants were the progeny of $bcd^{E1}$ $nos^{L7}$ $tsl^{146}$ homozygous females and expressed Resille:GFP to visualize cell outlines.

**Time-Lapse Imaging.** Embryos aged 2-4 hours were dechorionated for 2 min in 50% (vol/vol) bleach, washed in distilled water, and mounted in halocarbon oil 27 and 700, 1:1 (Sigma) between a coverslip and an oxygen-permeable membrane (YSI). The ventrolateral region of the embryo was imaged on a Zeiss LSM880 laser scanning confocal microscope with a 40X/1.2 NA water-immersion objective. Z-stacks were acquired at 1-µm steps and 15-s time intervals. Maximum intensity z-projections of 3 µm in the apical junctional plane were analyzed.

**Tissue Elongation Measurement.** Tissue elongation was measured by particle image velocimetry (PIV) using PIVlab in MATLAB (2). Each image was divided into 2-pass Fast-Fourier-Transform windows (120 × 120 pixels) with 50% overlaps. A displacement vector field for each window and each time point was determined by cross-correlating each window in the current time point and the image in the next time point. Tissue length change was measured by quantifying the cumulative sum of the anterior-directed displacement at the anterior end of the germband and the posterior-directed displacement at the posterior end of the germband. The onset of tissue elongation ($t = 0$) was the time point when the derivative of the tissue elongation curve intersects zero.

**Automated Image Segmentation and Cell Rearrangement Analysis.** Time-lapse movies were projected and despeckled using ImageJ. Processed movies were segmented and computationally analyzed using the MATLAB based software SEGGA, and errors were corrected manually with the interactive user interface (3). Cells were tracked and analyzed between $t = -10$ min and $t = 30$ min for each movie. To be included in cell rearrangement analysis, cells must be in the region of interest for at least 5 minutes after $t = 0$. The cell rearrangement rate shown is an equally weighted average over 1.5 minutes.

**Cell Shape Index and Cell Shape Alignment Analysis.** Based on the cell segmentation data, we computed the average cell shape index $\bar{p}$ by quantifying for each segmented cell both the perimeter $P$ and area $A$ of the polygon defined by the cell vertices (i.e. the points where at least 3 cells meet). The average cell shape index $\bar{p}$ at a time point is the average of $P/\sqrt{A}$ over all segmented cells. Cell shape alignment $Q$ was quantified using the triangle method (4–6). A triangular tiling was created based on the barycenters of the cellular polygons. For each triangle, we computed a symmetric, traceless shape tensor $\boldsymbol{q}$ quantifying triangle elongation (6). The area-weighted average of these tensors is the cell shape alignment tensor

$$\boldsymbol{Q} = \begin{pmatrix} Q_{xx} & Q_{xy} \\ Q_{xy} & -Q_{xx} \end{pmatrix} \quad \text{(S1)}$$

The cell shape alignment parameter $Q$ in the main text is the magnitude of this tensor defined by $Q = \left[Q_{xx}^2 + Q_{xy}^2\right]^{1/2}$.

**Vertex Model.** Our vertex model describes an epithelial tissue as a planar tiling of $N$ cellular polygons, where the degrees of freedom are the vertex positions $r_{k\alpha}$ (7). We use Latin indices starting with $k$ to refer to vertices and Greek indices starting with $\alpha$ to refer to spatial dimensions. Forces are defined such that cell perimeters and areas act as effective springs with a preferred perimeter $p_0$ and a preferred area of one. This is implemented via the following effective energy functional, which in dimensionless form is (8):

$$E = \sum_{i=1}^{N}[(p_i - p_0)^2 + k_A(a_i - 1)^2] \tag{S2}$$

Here, the sum is over all cells $i$, with perimeter $p_i$ and area $a_i$. The parameter $k_A$ is a dimensionless number comparing area and perimeter rigidity. We use periodic boundary conditions with box size $L_x \times L_y$ such that the average cell number density is one: $L_x L_y = N$. The boundary conditions can accommodate a skew (as in Lees-Edwards boundary conditions) with a corresponding simple shear $\gamma$. Hence, the system energy is a function of all vertex positions and the periodic box parameters: $E = E(\{r_{k\alpha}\}, L_x, L_y, \gamma)$. We focus on stable, force-balanced states of the system, which corresponds to local minima of $E$. To numerically find such states, we use the BFGS2 multidimensional minimization routine of the Gnu scientific library (GPL) with a cutoff on the average residual force of $10^{-6}$. We allow for manifold vertices – i.e. vertices are allowed to be in contact with more than three cells at once. During the minimization, the vertices belonging to an edge are fused to a single vertex whenever the edge length is below a cutoff of $10^{-3}$, and a vertex with at least four edges attached to it splits into several vertices whenever this is energetically favorable. While it is known that the existence of manifold vertices can change the transition point in vertex models (9), we checked that the energy-minimized states we obtained rarely contained any manifold vertices. In all simulations, we have $N = 512$ cells and $k_A = 1$.

For a given local energy minimum, we compute the simple shear modulus $G$ as described in (10):

$$G = \frac{1}{N}\left(\frac{\partial^2 E}{\partial \gamma^2} - \sum_m \frac{1}{\omega_m^2}\left[\sum_{k,\alpha} \frac{\partial^2 E}{\partial \gamma \partial r_{k\alpha}} u_{k\alpha}^m\right]^2\right) \tag{S3}$$

In the second term, the outer sum is over all positive eigenvalues $\omega_m^2$ and the corresponding eigenvectors $u_{k\alpha}^m$ of the Hessian matrix $(\partial^2 E/\partial r_{k\alpha}\partial r_{l\beta})$. In practice, we include all eigenvalues smaller than $10^{-14}$ in the sum. The inner sum in the second term is over all vertices and both spatial dimensions.

***Anisotropic vertex model.*** In all our simulations, we initialize the system with the Voronoi tessellation of a uniformly random point pattern on a squared domain ($L_x = L_y = L_0$). For the first set of simulations of anisotropic tissue, we apply an external pure shear strain $\varepsilon$ by setting $L_x = e^{\varepsilon} L_0$ and $L_y = e^{-\varepsilon} L_0$. We start with $\varepsilon = 0$ and increase in steps of 0.02 up to a value of $\varepsilon = 2$, minimizing the energy after each step. For these minimizations, we vary all vertex positions, keep the box dimensions fixed, but also allow the simple shear variable $\gamma$ to vary (shear-stabilized minimization). We follow this protocol for different values of $p_0$, which we varied between 3.5 and 4.5 in steps of 0.01. For each value of $p_0$ we carry out 100 separate simulation runs.

For the second set of simulations of an anisotropic tissue, we model the anisotropic myosin distribution in the germband by introducing an additional anisotropic line tension with amplitude $\lambda_0$ into the effective energy functional:

$$E = \sum_{i=1}^{N}[(p_i - p_0)^2 + k_A(a_i - 1)^2] + \sum_{\langle k,l\rangle} \lambda_{\langle k,l\rangle}\ell_{\langle k,l\rangle} \tag{S4}$$

While the first sum is the same as in Eq. (S2), we have added a second sum, which is over all edges in the system, connecting two vertices $k, l$. Here, $\ell_{\langle k,l\rangle}$ denotes the length the edge $\langle k, l\rangle$, and $\lambda_{\langle k,l\rangle}$ is a line tension associated with this edge. Before each minimization, we define each of these line tensions based on the respective edge angle $\theta_{\langle k,l\rangle}$ as follows:

$$\lambda_{\langle k,l\rangle} = \lambda_0 \cos(2[\theta_{\langle k,l\rangle} - \phi]) \tag{S5}$$

Thus, the line tension will be increased by $\lambda_0$ for edges parallel to lines with angle $\phi$ and decreased by $\lambda_0$ for edges perpendicular to that.

During each minimization run, we vary all vertex positions and the pure shear strain $\varepsilon = 1/2 \log(L_x/L_y)$, but keep the system area $L_xL_y$ and the simple shear strain $\gamma$ fixed. While we set $\lambda_{\langle k,l\rangle}$ before a minimization run and keep it constant during the minimization, the angle $\theta_{\langle k,l\rangle}$ usually changes during the minimization as the vertex positions are varied. As a consequence, the state obtained after the minimization will not correspond to an energy minimum anymore once we update the line tensions $\lambda_{\langle k,l\rangle}$ with the new angles $\theta_{\langle k,l\rangle}$. Thus, to identify a force-balanced state where the line tensions are consistent with the directions of the cell edges, we iterate over several minimizations, where after each minimization we update $\lambda_{\langle k,l\rangle}$ based on the latest angles $\theta_{\langle k,l\rangle}$. We stop these iterations once the states do not significantly change anymore, or more precisely, when the average residual stress per degree of freedom *before* a minimization, but with the *new* $\lambda_{\langle k,l\rangle}$, is smaller than $2 \times 10^{-6}$. We intentionally *do not* include the explicit dependency of $\lambda_{\langle k,l\rangle}$ on $\theta_{\langle k,l\rangle}$ and thus the vertex positions in our energy minimizations, because this would create additional torques in our model, while here we merely want to study the effect of an anisotropic distribution of line tensions as provided for instance by an anisotropic myosin distribution.

We set the direction of line tension anisotropy parallel to the $y$ axis, i.e. $\phi = \pi/2$ and varied the magnitude of line tension anisotropy $\lambda_0$ between zero and one in steps of 0.01. Again, the preferred perimeter $p_0$ is varied between 3.5 and 4.5 in steps of 0.01, where for each value of $p_0$ we run 100 separate simulations.

We found many states where the system flowed during a minimization until $\varepsilon$ was so large that the system was only one cell thick in the $y$ direction. In particular, this was the case in what was otherwise expected to be the floppy regime (cf. Fig. 3*B,C*). This will probably not only occur in the floppy regime, but also in the solid regime whenever the anisotropic stress created by the line tension anisotropy is large enough to overcome the yield stress, which perhaps explains why there is a gap between mechanically stable solid states and the black line in Fig. 3*C*. Because we did not obtain any force-balanced state of bulk vertex model tissue in this regime, we have no way to determine from our simulations neither the shear modulus, nor the morphological quantities $\bar{p}$ and $Q$, and thus this regime does not appear in Fig. 3*C*. To access this regime, one needs to include dynamics into the model, e.g. including a viscosity or a substrate friction.

To obtain Fig. 3*B,C*, we binned all of our energy-minimized configurations with respect to $\bar{p}$ and $Q$ (which were computed as described above), and then computed the fraction of floppy configurations within each bin. A configuration was defined floppy when its shear modulus $G$ was below a cutoff value of $10^{-5}$.

***Packing dependence of transition point.*** To study the packing-dependence of the transition point, we annealed the isotropic vertex model tissue at different temperatures prior to quenching the system to zero temperature, as this is a standard method for altering packing disorder in other materials such as structural glasses. To simulate the vertex model at a given temperature, we followed an Euler integration scheme updating all vertex positions $r_{k\alpha}$ as follows in each time step $\Delta t$:

$$r_{k\alpha} \rightarrow \; r_{k\alpha} + \mu F_{k\alpha} \Delta t + \eta_{k\alpha} \tag{S6}$$

Here, we have non-dimensionalized time such that the dimensionless motility $\mu$ is one, $F_{k\alpha} = -\partial E/\partial r_{k\alpha}$ is the force on vertex $k$ with the energy given by Eq. (S2), and $\eta_{k\alpha}$ is a normal distributed random force with zero average and variance $\langle \eta_{k\alpha} \eta_{l\beta} \rangle = 2\mu T \Delta t \delta_{kl} \delta_{\alpha\beta}$. To simulate these dynamics, we use the publicly available cellGPU code (11), with a time step of $\Delta t = 0.01$. For these simulations, vertices are always 3-fold coordinated and an edge undergoes a full T1 transition whenever its length is below a cutoff of 0.04.

We run 100 simulations for each set of parameters $(T, p_0)$, where $T$ varies logarithmically between $5 \times 10^{-6}$ and $1.5 \times 10^{-1}$, and $p_0$ varies between 3.7 and 3.9 in steps of 0.01. All simulations are thermalized at their target temperature for a time of $10^4$ before recording the data. We then perform simulations for $10^6$ at the target temperature before the temperature is quenched to $T = 0$. We calculate the decay of the self-overlap function for the slowest of the most solid states (low $p_0$ and $T$ sets) and confirm that the vertices are displaced less than a characteristic distance of 1/e at time $10^6$, suggesting the states are relaxed. To quench the temperature to zero, we run Eq. (S6) for an additional time of $10^6$. Afterwards, we use the BFGS2 algorithm of the GSL to further minimize until the average residual force per degree of freedom is below $10^{-6}$. The data are shown in Fig. S1, demonstrating that the transition point ${p_0}^*$ does depend systematically on the annealing temperature and therefore on the packing disorder.

The transition point we find occasionally decreases below the value of 3.81 (Fig. S1), which is the minimal transition point we would expect for disordered packings (12). To test whether this could be due to partial crystallization, we also quantified a hexatic order parameter:

$$\Phi_6 = \frac{1}{N_e} \sum_{\langle k,l \rangle} e^{6i\theta_{\langle k,l \rangle}} \tag{S7}$$

Here, the sum is over all $N_e$ edges in the system, where $\theta_{\langle k,l \rangle}$ is the angle of the edge between vertices $k$ and $l$. We find that the decreased transition point is indeed correlated with hexatic order $|\Phi_6|^2$ (Fig. S1).

**Theoretical Expectation for the Shift of the Transition Point.** In a recent publication (8), some of us showed that the transition point $\bar{p}_{\text{crit}}$ in the vertex model is expected to shift away from the isotropic transition point $p_o^*$ as the material is anisotropically deformed with strain $\varepsilon = 1/2 \log(L_x/L_y)$ as:

$$\bar{p}_{\text{crit}} = p_o^* + 4b\varepsilon^2 \tag{S8}$$

Here, $b$ is a constant prefactor whose precise value depends on the packing disorder, but whose typical value was previously found to be 0.6±0.2 (average ± standard deviation). We use here the pure shear strain variable $\varepsilon = 1/2 \log(L_x/L_y)$, which is related to the strain variable $\gamma$ used in

Ref. (8) as $\varepsilon = \gamma/2$, and so we get an additional factor of 4 in front of $b$ in Eq. (S8). However, Eq. (S8) has so far only been discussed without cell rearrangements, which do occur in our simulations.

To apply these ideas here, we replace the externally applied strain $\varepsilon$ with a kind of "internal" strain $\varepsilon_i$. To define this internal strain for some anisotropic vertex model configuration, we ask for the amount of strain $\varepsilon_i$ that is required to obtain this configuration from some virtual isotropic configuration without any cell rearrangements. Assuming that the transition point for this virtual isotropic state is the known $p_o^*$, we can then use Eq. (S8) with $\varepsilon_i$ to obtain the transition point for our anisotropic state. We choose to define tissue anisotropy by an order parameter $Q$ quantifying elongation and alignment of the cells (see above) (6). Thus, we are asking for the strain $\varepsilon_i$ needed to transform an isotropic state with $Q = 0$ to an anisotropic state with finite $Q$. In Ref. (6), some of us have shown that in the case of homogeneous deformation, this strain exactly corresponds to $\varepsilon_i = Q$ if we measure $Q$ based on a triangular tiling of the tissue. Hence, for tissue anisotropy $Q$ measured this way, we obtain:

$$\bar{p}_{\text{crit}} = p_o^* + 4bQ^2 \tag{S9}$$

An alternative way to obtain this equation is to Taylor expand $\bar{p}_{\text{crit}}$ in terms of the cell shape order tensor $\boldsymbol{Q}$, where the lowest-order term besides the constant allowed by symmetry is a term $\sim Q^2$. However, with the approach above, we can also connect the value of the prefactor $b$ to previous results.

Finally, the predictions in Ref. (8) strictly speaking refer to the non-dimensionalized average perimeter, i.e. the average of $p_i$ over all cells, whereas here by $\bar{p}$ we refer to the average shape index, i.e. the average of $p_i/\sqrt{a_i}$ over all cells. We verified that this difference does not play a role in our vertex model simulations.

***Fit to simulation data.*** To fit Eq. (S9) to the simulation data where we apply the external anisotropic deformation (Fig. 3*B*), we compute the average transition point for each $Q$ by interpreting the $\bar{p}$-dependent fraction of floppy configurations for fixed $Q$ as a cumulative probability distribution function and extracting the average from it. For varying $Q$, the resulting plot together with a fit to Eq. (S9) is shown in Fig. S2*A*.

We excluded a few data points from the fit, which were affected by the excess of rigid states observed around $\bar{p} \approx 4.15$ and $Q \approx 0.3$. It is so far not clear where this excess of rigid states comes from, but we speculate that it could be related to the system switching to a different kind of packing disorder in this region.

***Fits to experimental data.*** We fit Eq. (1) to the experimental data varying $p_0^*$ and keeping $b = 0.43$ constant, which is the value we found in the fit to the simulation data (Fig. S2*A*). We perform the fit such that the number of experimental data points on the wrong side of the fit curve, $n_{\text{tot}}$, is minimized. The number $n_{\text{tot}}$ is the sum of the number of solid data points above the fit curve and of fluid data points below the curve. To define tissue as solid, we use a cutoff on the cell rearrangement rate of 0.02 rearrangements per cell and minute. While different cutoffs lead to different fit values for $p_0^*$ (Fig. S2*B*), our main conclusions remain unchanged.

**Statistical Analysis.** Unless otherwise noted, error bars are the standard deviation.

**Data Availability.** Data and custom scripts are available from the corresponding author upon reasonable request.

### Supplementary Figures

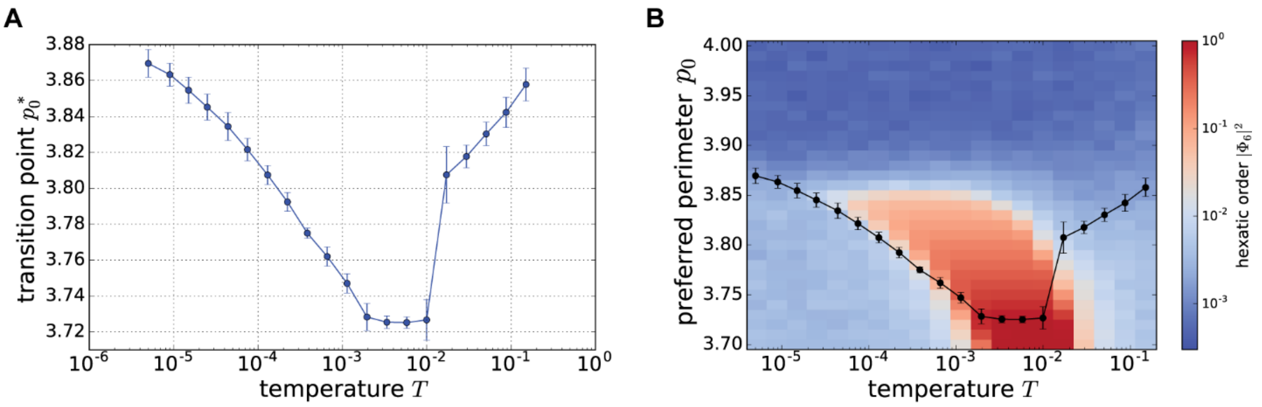


**Figure S1. The vertex model transition point depends on the packing disorder.** Results of vertex model simulations, where the effect of packing disorder is studied by annealing the model tissue with thermal fluctuations prior to quenching to a force-balanced state to create packings with different degrees of disorder. Taken together, these simulation results show that the critical shape index in the vertex model depends on the cellular packing disorder, where fluctuations can help to decrease packing disorder and $p_o^*$. (*A*) The transition point $p_o^*$ decreases with the annealing temperature $T$ before increasing again for very high $T$, confirming a dependence of the transition point on packing disorder. The values for $p_o^*$ decreased from 3.86 for low temperatures to 3.72 for higher temperatures, and increased again for even higher temperatures. (*B*) While the lower bound is below the previously determined limit of 3.81, this may be related to partial crystallization of the tissue in this regime. The hexatic bond-orientational order parameter, shown here depending on the preferred shape index $p_0$ and annealing temperature $T$, indicates at least partial crystallization for intermediate temperatures, which correlates with lower transition points $p_o^*$.

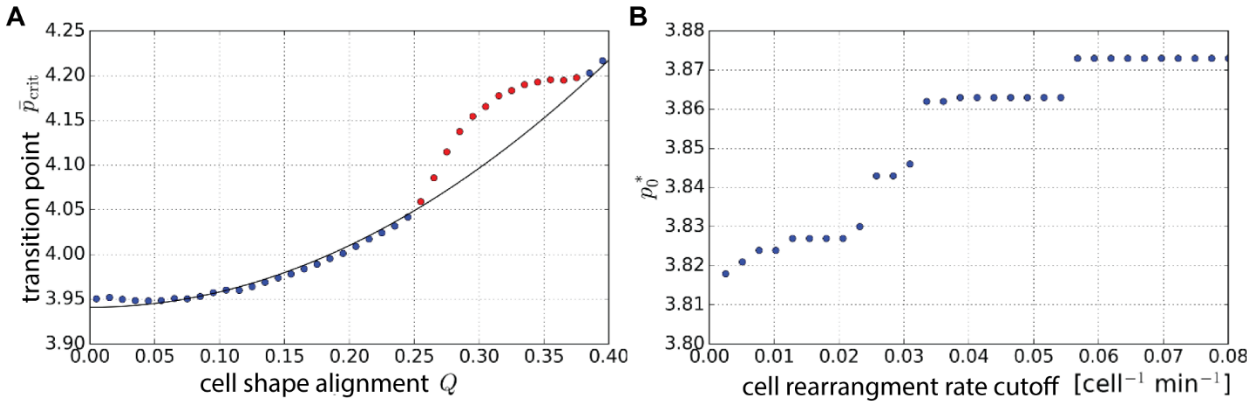


**Figure S2. Fits of Eq. (1) to simulations and experimental data.**
(*A*) Fit of Eq. (1) (black line) to the vertex model simulation results for the case of external deformation (dots, cf. Fig. 3*B*). The average transition points $\bar{p}_{\text{crit}}$ were extracted from the simulation data (Fig. 3*B*) by interpreting the fraction of floppy networks for fixed $Q$ as a cumulative probability density and extracting the average from it. From the fit we find $p_o^* = 3.94$ and $b = 0.43$. The red data points were excluded from the fit. These points are related to the excess number of rigid states around $Q \approx 0.3$ and $\bar{p} \approx 4.15$ (cf. Fig. 3*B*). (*B*) Dependence of the fit parameter $p_0^*$ on the cell rearrangement rate cutoff in the fit of Eq. (1) to the wild-type data (cf. Fig. 4*C*).
